# The mirtron miR-1010 functions in concert with its host gene SKIP to maintain synaptic homeostasis

**DOI:** 10.1101/353888

**Authors:** Christopher Amourda, Timothy E. Saunders

**Affiliations:** Mechanobiology Institute, National University of Singapore, Singapore; Department of Biological Sciences, National University of Singapore, Singapore; Institute of Molecular and Cell Biology, A*Star, Proteos, Singapore; Lead Contact

**Keywords:** mirtron function, miR-1010, synaptic homeostasis, nicotinic receptor, Adf-1, incoherent feedforward loop, negative feedback loop

## Abstract

Mirtrons are non-canonical miRNAs arising by splicing and debranching from short introns. A plethora of introns have been inferred by computational analyses as potential mirtrons. Yet, few have been experimentally validated and their functions, particularly in relation to their host genes, remain poorly understood. Here, we found that larvae lacking the mirtron miR-1010 are unable to grow properly and pupariate. We show that miR-1010 downregulates nAcRβ2. Increase of cortical nAcRβ2 mediated by neural activity elevates the level of intracellular Ca^2+^, which in turn activates CaMKII and, further downstream, the transcription factor Adf-1. We reveal that Adf-1 initiates the expression of SKIP, the host gene of miR-1010. Preventing synaptic potentials from overshooting their optimal range requires both SKIP to temper synaptic potentials (incoherent feedforward loop) and miR-1010 to reduce nAcRβ2 mRNA levels (negative feedback loop). Our results demonstrate how a mirtron, in coordination with its host gene, contributes to maintaining homeostasis.

## Introduction

Since their discovery more than two decades ago (Lee, Feinbaum and Ambros, 1993; Wightman, Ha and Ruvkun, 1993), microRNAs (miRNAs) have been established as key cellular micromanagers. MiRNAs control nearly all cellular pathways, with a large subset being indispensable to development (Lucchetta, Carthew and Ismagilov, 2009; Carthew, Agbu and Giri, 2017; Mok *et al.*, 2018). Canonical miRNAs are genetically encoded short (18-22 nucleotides) hairpin structures that are recognised, after transcription, by the RNAse type III endonuclease Drosha which cleaves the stem region in the nucleus (Denli, A.M., Tops, B.B.J., Plasterk, R.H.A., Ketting, R.F., Hannon, 2004; Gregory *et al.*, 2004; Berezikov, 2011). MiRNAs are subsequently exported from the nucleus and undergo a final step of maturation before being functionally able to regulate the mRNA level of their target genes (Pasquinelli, Hunter and Bracht, 2005; Giri and Carthew, 2014; Carthew, Agbu and Giri, 2017; Chandra *et al.*, 2017). Recent computational and experimental efforts identified a non-canonical miRNA maturation pathway in which introns are debranched from precursor mRNAs by the Lariat debranching enzyme and enter the miRNA processing pathway without requiring cleavage by Drosha (Okamura *et al.*, 2007; Ruby, Jan and Bartel, 2007; Westholm and Lai, 2011). This new class of miRNAs were named mirtrons and numerous introns have, subsequently, been inferred as potential mirtrons (Chung *et al.*, 2011; Ladewig *et al.*, 2012; Wen *et al.*, 2015). Further studies revealed that mirtrons are widespread across taxa and some show a high degree of conservation (Curtis, Sibley and Wood, 2012), implying important regulatory functions.

The repertoire of canonical miRNA genomic sources is characterised by its versatility. Indeed, miRNAs can be found as single or clustered transcriptional units bearing their own regulatory elements. MiRNAs are also found within introns of host genes both in sense or anti-sense orientations, indicating that their expression does not necessarily correlates with that of their host genes (Rodriguez *et al.*, 2004; Martinez *et al.*, 2008; Ozsolak *et al.*, 2008; Isik and Berezikov, 2013). An important question arising from the genomic organisation of canonical miRNAs is: what is the reason for the emergence of mirtrons? It has been hypothesised that alternative miRNA processing pathways may be important in stressful conditions. Anaerobic conditions in tumors or exposure to hormones lead to a down-regulation or inhibition of the miRNA processing components (Merritt *et al.*, 2008; Faggad *et al.*, 2010). For example, Drosha mRNA level is reduced by around 50% in ovarian-cancer specimens. In such circumstances, pathways regulated by miRNAs are perturbed (Merritt *et al.*, 2008). Although, the presence of mirtrons in such stressful conditions has not been documented, an attractive possibility is that mirtrons fulfil fundamental roles under stressful conditions to maintain cellular homeostasis (Curtis, Sibley and Wood, 2012). In particular, since mirtrons mature regardless of the level of Drosha, their processing and maturation should not be affected under stress as the transcription machinery remains functional. Alternatively, mirtrons may have emerged from mutation of short intronic sequences that evolved into hairpin structures (Berezikov *et al.*, 2007; Curtis, Sibley and Wood, 2012). Importantly, the biological significance of intronic miRNAs and, especially, mirtrons must be understood with respect to the function of their host gene (Inui *et al.*, 2018).

Here, we show that the mirtron miR-1010 regulates the level of the nicotinic acetylcholine receptor b2 (nAcRβ2) via a negative feedback loop whereby elevated nAcRβ2 results in increased miR-1010 levels. In the absence of miR-1010 larvae cease growth and do not pupariate. Elevated levels of nAcRβ2 upon neural activity also triggers a homeostatic response whereby SKIP, the host gene of miR-1010, amplifies the Shal K^+^ channel role in tempering membrane potentials (Diao, Waro and Tsunoda, 2009; Ping and Tsunoda, 2011). However, this negative feedforward response is dispensable, with viable adults emerging in SKIP mutants. Our work demonstrates that miR-1010, transcribed alongside SKIP, is involved in a critical negative feedback loop that decreases nAcRβ2 mRNA levels to restore homeostasis. Importantly, we deconstruct the underlying mechanisms of the feedforward and feedback loops that regulate the neuron potential, and we find that the presence of miR-1010 is vital in this process, in contrast to that of SKIP. Finally, we show that miR-1010 is upregulated upon exposure to nicotine. Therefore, we believe that our results will be contribute to further our understanding of nicotine-related disorders.

## Results and Discussion

### MiR-1010, in contrast to its host gene SKIP, is indispensable to viability

We brought our attention to the mirtron miR-1010 as homozygous mutants are lethal (Chen *et al.*, 2014). MiR-1010 is located within the Shal potassium (K^+^) channel Interacting Protein (SKIP) gene, between exons 4 and 5 (Figures. 1A and S1). Replacing miR-1010 by a LoxP sequence does not affect embryogenesis as larvae hatch without any apparent defects and with similar viability as wild-type animals. However, mutant larvae are unable to grow; they retain a first instar larva size (<0.5mg) and fail to pupariate even after ten days of larval life (Figure. 1B). The ecdysone pathway, a steroid hormone controlling larval moulting (Baehrecke, 1996; Yamanaka, Rewitz and O’Connor, 2013), appears unaffected with the Ecdysone-induced protein 75B (E75B), the Prothoracicotropic hormone (PTTH), Phantom (Phm) and Disembodied (Dib) being expressed within wild-type ranges (Figures S2A-E). Furthermore, the insulin receptor (dInr) and the *Drosophila* insulin-like peptide 2 (Dilp2) are upregulated to counteract the lack of growth (Figures S2F and S2G). The standard level of chico (substrate of the insulin receptor (Boulan, Milán and Léoplold, 2016)), sNPF and ns3 (both involved in dilp2 secretion (Boulan, Milán and Léoplold, 2016)) concur with the insulin pathway and dilp2 secretion being operative in miR-1010^-/-^ (Figures S2H-J). Moreover, we observed no defects in food intake in miR-1010^-/-^ larvae (Figure 1C). Strikingly, the metabolic brake 4E-BP (Teleman, Chen and Cohen, 2005) shows a 10-fold increase directly at the onset of larval life and remains at a high level in miR-1010 mutant larvae as compared to wild-type larvae (Figure 1D). The high level of 4E-BP indicates that larvae sense a stressful condition and, as a result, growth is inhibited until returning to more adequate conditions (Teleman, Chen and Cohen, 2005). Prior to investigating the causes underlying the miR-1010^-/-^ phenotype, we first sought to verify that the impeded growth is solely due to the lack of miR-1010 and not a result of an alteration of SKIP. For this purpose, we used a SKIP MiMIC (SKIP^-/-^) mutant where a triple stop codon is inserted within the first intron (Venken *et al.*, 2011). Importantly, the MiMIC insertion does not prevent transcription but alters the translation of a given gene. Hence, the miR-1010 level remains at a wild-type level in SKIP^-/-^ larvae (Figure S2K). Concordantly, SKIP^-/-^ larvae show a mild timing phenotype and pupate with a 24h delay (Figure 1B). These results indicate that the deficit in larval growth is due to a lack of miR-1010.

**Figure 1.**
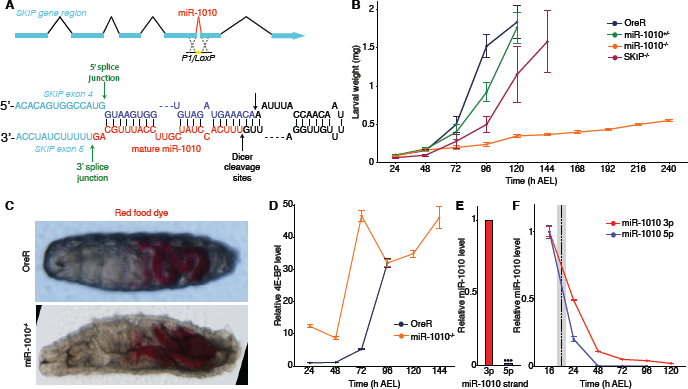
Larvae fail to grow and pupariate in miR-1010^-/-^. (**A**) The mirtron miR-1010 is located in the SKIP gene, between the exons 4 and 5. The entire miR-1010 has been replaced by a LoxP sequence in the miR-1010^-/-^ line (top panel). MiR-1010 forms a hairpin loop and is debranched from SKIP by the splicing machinery and further matures as a normal miRNA (bottom panel). (**B**) Larval weight was measured directly after hatching until pupariation. OreR, miR-1010^+/-^ and SKIP^-/-^ reached ˜2mg before pupariation. MiR-1010^-/-^ weight was recorded for 10 days but did not pupariate. (**C**) Stereoscope pictures of OreR and miR-1010^-/-^ larvae allowed to eat coloured yeast paste show no feeding behaviour defect. (**D**) Larval 4E-BP transcript levels measured by RT-qPCR in OreR and miR-1010^-/-^. Fold changes are relative to OreR at 24h AEL. (**E**-**F**) MiR-1010 strand specific RT-qPCR shows a strong bias toward miR-1010 3p (**E**) and a high expression during embryogenesis before gradually decreasing during larval stages (**F**). Expression level are relative to the expression at 16h AEL. The dashed line indicates the average hatching time and the grey rectangle represents the standard deviation in hatching times at 25ºC. All values are means ± SD (***P<0.001, n = at least 9 for each experiment).

### nAcRβ2 is the main target of miR-1010

We next sought to identify the targets of miR-1010. In contrast to canonical miRNAs, mirtrons 3’ arms are preferentially more stable than their 5’ complement (Okamura *et al.*, 2007). Consistent with this biased stability, we found only traces of miR-1010 5’ strand throughout development (Figure 1E). Further, we observed that miR-1010 has its highest expression during mid-embryogenesis (16h AEL) and gradually decreases during larval development (Figure 1F). We next used a Gal4 driver inserted downstream of the SKIP regulatory region (SKIP>Gal4) to locate the spatial expression of both SKIP and miR-1010. We assume here that SKIP and miR-1010 have the same expression pattern as miR-1010 arises from the debranching of SKIP introns. We found that SKIP and miR-1010 are expressed in the central nervous system (CNS) and more prominently in the axons emanating from the CNS (Figure 2A and Movie 1). Coupling this expression pattern with the computationally predicted targets (Lewis, Burge and Bartel, 2005; Kheradpour *et al.*, 2007; Ruby *et al.*, 2007; Ruby, Jan and Bartel, 2007) for the miR-1010 3’ strand, we refined a list of targets and examined their expression in miR-1010^-/-^. Amongst 36 tested targets, we found that Drl-2, nAcRβ2 and CG3078 are (i) expressed at meaningful level (judged by the Ct values in qPCR experiments) and (ii) consistently overexpressed (generally >2-fold for all targets) in miR-1010^-/-^ throughout larval development (Figures 2B, 2D and S3a). Of these, only nAcRβ2 shows higher expression during embryogenesis (Figure S3B). Notably, the nAcRβ2 is amongst the most prominently expressed nAcRs during early larval life and further the only subtype overexpressed in miR-1010^-/-^ (Figures S3C and S3D). We infer from these data that nAcRβ2 plays a crucial role at this stage of development and accounts, in part, for the phenotype observed in miR-1010^-/-^. Alongside the predicted targets, we noticed that the mRNA levels of SKIP and Shal in miR-1010^-/-^ larvae show a ˜3-fold increase as compared to control larvae (Figure 2E). The increase in mRNA level translates into an increase in protein level for both nAcRβ2 and SKIP (Figures S3E and S3F). Importantly, the nAcRβ2 protein level is slightly increased during embryogenesis but dramatically greater at larval stages in miR-1010^-/-^ (Figure S3E). SKIP has two shorter isoforms besides its full length protein (Diao, Waro and Tsunoda, 2009) and we observed an upregulation of both the full length and the SKIP3 isoform at larval stages (Figure S3F). SKIP3 has previously been shown to favour the slow inactivation mode of Shal (Diao, Waro and Tsunoda, 2009). These results suggest that miR-1010, its host gene, and targets are within the same regulatory pathway.

**Figure 2.**
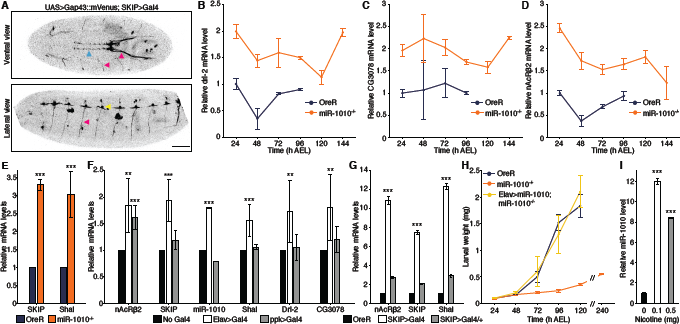
nAcRα2, Drl-2 and CG3078 levels is elevated in miR-1010^-/-^. (**A**) Confocal imaging of embryos expressing a membrane marker (UAS>Gap43::mVenus) driven by SKIP>Gal4. SKIP/miR-1010 are expressed in the CNS (blue arrowhead), in axons (pink arrowheads) emanating from the CNS and at neuromuscular junctions (yellow arrowhead). Scale bar is 50µm. (**B**-**D**) Drl-2 (**B**), CG3078 (**C**) and nAcRβ2 (**D**) transcript levels (RT-qPCR) in miR-1010^-/-^ (orange). Fold changes are relative to OreR (dark blue) at 24h AEL. (**E**) SKIP and Shal transcript levels (RT-qPCR) in miR-1010^-/-^ relative to OreR larvae at 24h AEL. (**F**) Transcripts levels (RT-qPCR) in nAcRβ2 overexpressed by Elav>Gal4 (white) or ppk>Gal4 (grey) relative to non-induce UAS>nAcRβ2 (black) at 24h AEL. (**G**) nAcRβ2, SKIP and Shal transcript levels (RT-qPCR) in homozygous SKIP>Gal4 (white) and in SKIP>Gal4/+ (gray) relative to OreR (black) at 24h AEL. (**H**) Larval weight was measured in OreR, miR-1010^-/-^ and in Elav>Gal4/UAS>miR-1010; miR-1010^-/-^ rescue line. (**i**) miR-1010 levels upon exposure to 0.1mg (white) and 0.5mg (gray) of nicotine as compared to 0mg (black). All values are means ± SD (*P<0.05, **P<0.01, ***P<0.001, n = at least 9 for each experiment).

### Overexpression of nAcRβ2 results in miR-1010 upregulation

We next attempted to rescue the miR-1010^-/-^ phenotype by knocking down specific targets. Mutants for Drl-2 and CG3078 fail to rescue the miR-1010^-/-^ phenotype (Figure S4A). We were unable to perform similar rescue experiment for the nAcRβ2 as the cytogenic locations of miR-1010 and nAcRβ2 are in close proximity (94A4 and 96A5, respectively) and therefore prevent chromosomal recombination. MiRNA deficiency essentially results in overexpression of its targets. We reasoned that overexpressing the coding sequence (i.e. without the 3’UTR bearing miR-1010 binding sites) of miR-1010 targets should phenocopy miR-1010^-/-^ (Figure S4B). We did not impede larval growth as in miR-1010^-/-^ (Figures S4C-S4E), nevertheless, pan-neural (Elav>Gal4) overexpression of nAcRβ2 resulted in an interesting pattern whereby SKIP, miR-1010 and Shal are upregulated along with Drl-2 and CG3078 (Figure 2F). On the contrary, overexpression of nAcRβ2 in sensory neurons using Pickpocket Gal4 driver (Adams *et al.*, 1998; Ainsley *et al.*, 2008) (ppk>Gal4) did not increase the level of SKIP, miR-1010 and Shal (Figure 2F). These results indicate that miR-1010 is neuron specific. Overexpressing Drl-2 or CG3078 with Elav>Gal4 does not affect the levels of nAcRβ2, SKIP and Shal (Figures S4F-S4G). Further, we noticed that the SKIP>Gal4 driver line (when homozygous) exhibits a high level of nAcRβ2 accompanied by high levels of SKIP and Shal (Figure 2G). The SKIP>Gal4 line is, however, homozygous viable. Therefore, this expression pattern indicates that SKIP transcription or introns debranching functions at a sub-optimal level in the SKIP>Gal4 line. This phenotype can be partially rescued upon restoring a copy of a wild-type chromosome (Figure 2G). Last, we were able to rescue the larval growth defects in miR-1010^-/-^ by expressing miR-1010 under the pan-neural driver Elav (Figures 2H and S4H-S4J). The mRNA levels of nAcRβ2, Shal, Drl-2 and CG3078 are brought down by the overexpression of miR-1010. However, the SKIP mRNA level is consistently higher (~10-fold) in the rescue line. This expression pattern is surprising and further experiments will be required to understand the reason underlying SKIP upregulation. Altogether, these experiments indicate that an increase in nAcRβ2 gives rise to an upregulation of SKIP, miR-1010 and Shal genes, a necessity to maintain the neural circuit within acceptable conditions. The upregulation of Drl-2 and CG3078 upon nAcRβ2 increase is intriguing. The most likely explanation is that the higher levels of Drl-2 and CG3078 in miR-1010^-/-^ larvae is simply a response to increased neural activity, not the direct result of a lack of miR-1010. CG3078 upregulation upon neural activity is plausible as its inferred function is to regulate K^+^ channel activity (Gramates *et al.*, 2017). However, the reason for Drl-2, an axon guiding protein (Grillenzoni *et al.*, 2007; Sakurai *et al.*, 2009; Reynaud *et al.*, 2015), upregulation is more enigmatic and may be a response to axonal growth following increased activity.

### Nicotine exposure elevates the level miR-1010

nAcRs boost synaptic potentials and trigger the upregulation of cortical nAcRs (Ping and Tsunoda, 2011). These receptors are permeable to Ca^2+^ and the subsequent influx of Ca^2+^ activates the CaMKII. In turn, CaMKII triggers the expression of the Shal K^+^ channel. Therefore, K^+^ ions are released from neurons and temper membrane potentials. Shal and its interacting protein SKIP have been shown to be important members of a pathway required to stabilise these synaptic potentials (Diao, Waro and Tsunoda, 2009; Ping and Tsunoda, 2011). Despite the apparent importance of such a pathway, we found that neither knockout of Shal nor SKIP resulted in lethality (Figure S4K). Combining these results, we hypothesised that the mRNA level of nAcRβ2 is downregulated by miR-1010 and that this negative feedback works complementarily to SKIP and Shal to temper synaptic potentials. As previously described, SKIP is presumed to modulate Shal channels inactivation kinetics by favouring Shal slow inactivation mode (Tsunoda and Salkoff, 1995; Diao, Waro and Tsunoda, 2009; Ping *et al.*, 2011). Crucially, however, our results suggest that the SKIP/Shal part of the pathway is only acting to temper the potential response and is not fundamental in restoring homeostasis. In contrast, the negative feedback loop mediated via miR-1010 is indispensable for returning the system to homeostasis after stimulation.

To further test this model, we exposed wild-type larvae to different doses (0.1 and 0.5mg) of nicotine, agonist of nAcRs, for 24h. We saw a striking ~10-fold increase in miR-1010 levels for both dosage conditions (Figure 2I). This result shows that there exists a direct relationship between nAcRs activation and miR-1010 expression. Importantly, it suggests that miR-1010 plays a hitherto underappreciated role in regulating synaptic potentials under stress conditions.

## Adf-1 controls SKIP, miR-1010 and Shal expression

We next examined the regulation of SKIP and Shal expression. Through computational analysis, we found that the Alcohol dehydrogenase transcription factor 1 (Adf-1) has predicted binding sequences in both Shal and SKIP regulatory regions (Messeguer *et al.*, 2002; Farré *et al.*, 2003) (Figure S5A). Further, Adf-1 is phosphorylated by CaMKII and is therefore a plausible candidate (Timmerman *et al.*, 2013). It has also been demonstrated by ChIP-seq experiments that Adf-1 is phosphorylated upon neural activity and blocks Fas2 and (indirectly) Staufen to allow neuronal growth (Timmerman *et al.*, 2013). We obtained the ChIP-seq results mentioned above and noticed that Adf-1 also binds to Shal and to a smaller extent to SKIP (Figure S5B). We confirmed these results by performing a ChIP-qPCR for Adf-1 and showed that Adf-1 strongly binds Fas2, Shal and SKIP regulatory regions in OreR and miR-1010^+/^- mutants. However, this binding is significantly reduced in miR-1010^-/-^ (Figure 3A). Adf-1 has been shown to positively correlate with high Pol II-pausing indices (Timmerman *et al.*, 2013). Therefore, we reasoned that in its non-phosphorylated state Adf-1 is bound to Shal and SKIP regulatory elements and prevents their transcription. Upon neural activity, Adf-1 is phosphorylated and releases Shal and SKIP expression. If our hypothesis is correct, we should then observe higher levels of SKIP, miR-1010 and Shal in Adf-1^-/-^ larvae. The majority of Adf-1^-/-^ (~80%) fail to complete embryogenesis (Dezazzo *et al.*, 2000). Those that survive to larval stages do not grow noticeably and display a phenotype reminiscent to that of miR-1010^-/-^ larvae (Figure 3B). We performed qPCR on Adf-1^-/-^ that reached the larval stage and we saw a general increase in mRNA levels for all candidates (Figure 3C). However, our results are quite variable, a phenomenon unsurprising given the pleiotropic function of Adf-1 (Dezazzo *et al.*, 2000; Timmerman *et al.*, 2013). Adf-1 promotes neuronal growth upon neural activity by shutting down Fas2 and Staufen. Mechanism(s) must be in place to prevent detrimental overgrowth. By having Adf-1 also regulate Shal and SKIP, the system couples growth to neural activity. This mechanism introduces an incoherent feedforward loop to ensure robust growth in response to neural activity (Figure 3D).

**Figure 3.**
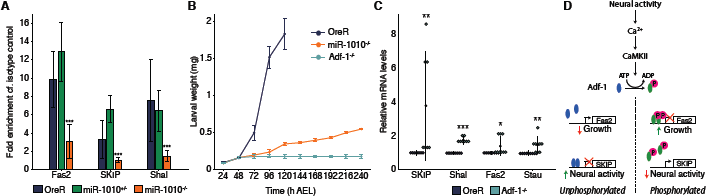
Adf-1 controls the expression of SKIP, miR-1010 and Shal. (**A**) ChIP-qPCR for Adf-1 performed in OreR (darkblue), miR-1010^+/^- (green) and miR-1010^-/-^ (orange). Fold enrichments are relative to the Rabbit IgG isotype control. (**B**) Larval weight in Adf-1^-/-^ as compared to OreR and miR-1010^-/-^. (**C**) Dot plot representing transcript levels measured in Adf-1^-/-^ (turquoise) relative to OreR (dark blue) at 24h AEL. Black dots and lines represent, respectively, the mean and the standard deviation for each condition. (**D**) Model of Adf-1-mediated coupling of neural activity and growth. Adf-1 is represented in blue in its nonphosphorylated form and in green upon phosphorylation. All values are means ± SD (*P<0.05, **P<0.01, ***P<0.001, n = at least 9 for each experiments).

### MiR-1010, in concert, with SKIP is necessary to maintain synaptic homeostasis

We have revealed here that nAcRβ2, Shal, SKIP and miR-1010 form a network that contains an incoherent feedforward loop coupled with a negative feedback loop (Figure 4A). Shal and SKIP are necessary to counteract the influx of positively charged ions upon neural activity. Nevertheless, Shal and SKIP are not sufficient to maintain membrane potentials within an optimal range and the regulation of nAcRβ2 by miR-1010 appears to be necessary to preserve synaptic homeostasis. Therefore, the mirtron miR-1010 and its host gene SKIP appear to work in tandem to ensure synaptic homeostasis (Figure 4B). To test this conclusion, we used mass-action kinetics to model the average long-term voltage potential response due to the two interaction loops outlined in Figure 4A after activation of nAcRβ2 (see Supplementary Note for details). We see that loss of SKIP or partial loss of miR-1010 results in delayed return of the potential to homeostatic levels (Figure 4C), consistent with our observed delay in developmental time (Figure 1B). However, complete loss of miR-1010 results in the potential staying active over a long period (Figure 4C), consistent with the observed stress response seen in miR-1010^-/-^. The modelling reveals that miR-1010 is effectively acting like a switch, controlling whether nAcRβ2 has high or low expression, and this switch-like behaviour is dependent on miR-1010 expression levels (see Supplementary Note for further discussion). The model predicts that the average membrane potential response will also be higher in miR-1010^+/^- and SKIP^-/-^ and it will be interesting to test this, for example using a GCaMP reporter.

**Figure 4.**
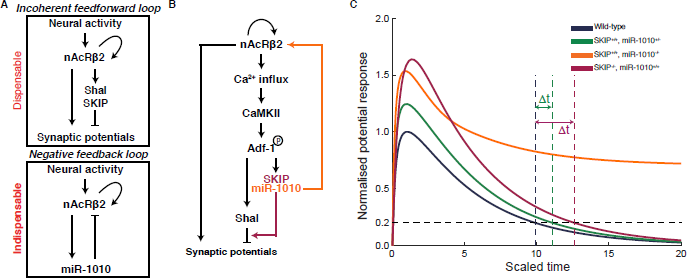
miR-1010 controls synaptic homeostasis by downregulating nAcRα2. (**A**) nAcRβ2 is part of incoherent feedforward loop to temper synaptic potentials and a negative feedback loop through which miR-1010 downregulates nAcRβ2 upon receptor activation. (**B**) Both loops work cooperatively to prevent synaptic potentials from overshooting their optimal range. (**C**) Long-term average potential response obtained by mass-action kinetic simulation of model outlined in (**A**). Resting potential defined to 0 (corresponding to ~-70meV). Curves normalised to maximum potential in wild-type simulation (parameters and description in Supplementary Note). Time is scaled relative to the lifetime of nAcRβ2. Δt represents delay in potential in miR-1010^+/^- and SKIP^-/-^ to return to less than 20% of peak response in wild-type conditions.

We have studied the role of miR-1010 in a developmental context but it is important to note that exposure to nicotine results in a dramatic increase of miR-1010 level in wild-type larvae. Hence, this pathway is likely not restricted to development and applies to later stages of life. Deciphering the relevance of miR-1010 regulation in nicotine-related disorder (such as nicotine addiction or Alzheimer disease) would constitute exciting avenues of research. Our work sheds light on the coherence between the functions of mirtrons and their host genes. In this case, SKIP and miR-1010 act in a coordinated fashion to efficiently maintain homeostasis. We predict that (i) such regulatory loops are not restricted to synaptic homeostasis and (ii) homologous pathways exist in higher organisms. MiR-1010 shares sequence homologies with the mammalian miR-412 (Ibáñez-Ventoso, Vora and Driscoll, 2008). However, miR-412 is not a mirtron and does not seem to directly regulate nAcRs (from computationally predicted targets). Rather, we believe that identifying a functional homolog would help to further understand the pertinence of homeostasis control by mirtrons.

## Acknowledgments

We thank Victor Corces and Stephen Cohen for sharing precious reagents and data. We acknowledge Katsutomo Okamura and all Saunders’ lab members for fruitful discussions. This work was supported by the National Research Foundation Singapore under an NRF Fellowship to T.E.S. (NRF2012NRF-NRFF001-094).

## Author Contributions

C.A. designed the study, performed the experiments, analysed the data and wrote the manuscript. T.E.S. performed the mathematical modelling, aided in the study design and statistical analysis, and contributed to writing the manuscript. Both authors agree to the final version of the manuscript.

The authors declare no competing financial interest.

